# Genome-wide assessment of genetic risk for systemic lupus erythematosus and disease severity

**DOI:** 10.1101/614867

**Authors:** Lingyan Chen, Yong-Fei Wang, Lu Liu, Adrianna Bielowka, Rahell Ahmed, Huoru Zhang, Phil Tombleson, Amy L Roberts, Christopher A Odhams, Deborah S Cunninghame Graham, Xuejun Zhang, Wangling Yang, Timothy J Vyse, David L Morris

## Abstract

**Objective:** Using three European and two Chinese genome-wide association studies (GWAS), we investigated the performance of genetic risk scores (GRS) for predicting the susceptibility and severity of Systemic lupus erythematosus (SLE), using renal disease as a proxy for severity.

**Methods:** We used four GWASs to test the performance of GRS both cross validating within the European population and between European and Chinese populations. The performance of GRS in SLE risk prediction was evaluated by Receiver Operating Characteristic (ROC) curves. We then analyzed the polygenic nature of SLE statistically. We also partitioned patients according to their age-of-onset and evaluated the predictability of GRS in disease severity in each age group.

**Results:** We found consistently that the best GRS in the prediction of SLE used SNPs associated at the level of *P*<1e-05 in all GWAS datasets and that SNPs with *P*-values above 0.2 were inflated for SLE true positive signals. The GRS results in an area under the ROC curve ranging between 0.64 and 0.72, within European and between the European and Chinese populations. We further showed a significant positive correlation between a GRS and renal disease in two independent European GWAS (*P_cohort1_*=2.44e-08; *P_cohort2_*=0.00205) and a significant negative correlation with age of SLE onset (*P_cohort1_*=1.76e-12; *P_cohort2_*=0.00384). We found that the GRS performed better in prediction of renal disease in the ‘later onset’ compared to the ‘earlier onset’ group.

**Conclusion:** The GRS predicts SLE in both European and Chinese populations and correlates with poorer prognostic factors: young age of onset and lupus nephritis.

## Introduction

Systemic lupus erythematosus (SLE [MIM: 601744]) is a chronic inflammatory autoimmune disease characterized by a wide spectrum of signs and symptoms varying among affected individuals and can involve many organs and systems, including the skin, joints, kidneys, lungs, central nervous system, and haematopoietic system (1). A recent report underscores that SLE is among the leading causes of death in young females, particular females among ages 15-24 years, in which SLE ranked tenth in the leading causes of death in all populations and fifth for African American and Hispanic females (2). Lupus nephritis is the most common cause of morbidity and mortality. Patients with kidney disease are likely to have more severe clinical outcomes and a shorter lifespan. 30-60% of adults and up to 70% of children with SLE have renal disease, characterized by the glomerular deposition of immune complexes and an ensuring inflammatory response (3). Genetic ancestry influences the incidence and prevalence of SLE and kidney involvement, being more frequent in Hispanics, Africans and Asians than in European (4–7). Currently, kidney disease in SLE is diagnosed by use of light microscopy, which drives therapeutic decisionmaking. However, not all patients will respond to therapy, indicating that additional information focusing on the mechanism of tissue injury is required. Moreover, early detection of kidney involvement in SLE is important because early treatment can be applied to reduce the accumulation of renal disability.

Although the exact aetiology of lupus is not fully understood, a strong genetic link has been identified through the application of family (8, 9) and twins studies (10). SLE does not follow a single locus Mendelian pattern of inheritance, and so it is termed a complex trait. Complex traits are multi-factorial with both genetic and environmental contributions. Genome-wide association studies (GWAS) have been successfully used to investigate the genetic basis of a disease and this has dramatically advanced knowledge of the genetic aetiology of SLE. Our recent review summarized a total of 84 genetic loci that are implicated as SLE risk (11). Despite the advances in the genetics of SLE, it is not clear how to utilise genetic information for the prediction of SLE risk or severity.

A genetic risk score (GRS) summarizes risk-associated variations by aggregating information from multiple risk single nucleotide polymorphisms (SNPs). The approach to calculate the GRS is to simply count disease-associated alleles or weighting the summed alleles by log Odds Ratios. Recent studies (12, 13) have proposed methods which select SNPs from GWAS by LD (linkage disequilibrium) pruning and clumping and thresholding for GRS calculation. As the number of SNPs included in a GRS increases, the distribution approaches normality, even when individual risk alleles are relatively uncommon. Therefore, a GRS can be an effective means of constructing a genome-wide risk measurement that summarises an individual’s genetic predisposition to SLE. Moreover, as GRSs pool information from multiple SNPs, each individual SNP does not strongly influence the summary measurement. Thus, the GRS is more robust to imperfect linkage for any tag SNP and causal SNP, and less sensitive to minor allele frequencies for individual SNPs (14–17).

Several studies (18–23) have looked at GRS for SLE, however many relied on very few SNPs (23), had sample sizes inadequate for GRS, did not compare results across populations or were restricted to SNPs on the Immunochip. We investigated, for the first time, the performance of genome-wide SNPs for predicting SLE. As in the most recent study of Lupus Nephritis (21) we also investigated the predictive performance of SNPs published as associated with SLE for disease severity. This study used data on three European GWAS and two Chinese GWAS. We first tested whether a quantitative model - a GRS derived from SLE GWAS applying a range of methods using genome wide SNPs, was an effective way to distinguish SLE patients and controls in three independent European cohorts. Next, we classified SLE patients into two groups: SLE renal+ (patients with renal disease) and SLE renal- (patients without renal disease), and performed a case-case genome-wide association study (GWAS) in two independent SLE cohorts with available renal data for the identification of SLE renal susceptibility loci. We then tested whether a GRS derived from SLE GWAS was an effective way to distinguish SLE patients with or without renal disease in two independent cohorts. A GRS analysis for SLE was performed across Chinese and European data where we trained the GRS in one population and predicted in the other. The SLE risk score was elevated in those with renal disease (compared to those without) and it showed a negative correlation with age of onset of the disease.

## Patients and Methods

### Samples source

European samples were from three previously published SLE GWAS – the SLE main cohort (24), the SLEGEN cohort (25), and the Genentech cohort (26). The SLE main cohort (24) was the biggest SLE GWAS, which consisted of 4,036 SLE patients and 6,959 healthy controls. A total number of 603,208 SNPs were available post quality control. The SLEGEN cohort (25) was carried out by The International Consortium for Systemic Lupus Erythematosus Genetics (SLEGEN) on women of European ancestry, which comprised 283,211 SNPs genotyped for 2,542 controls and 533 SLE patients. The Genentech cohort (26) was performed by Genentech on North American individuals of European descent, which comprised 487,208 SNPs genotyped for 1,165 cases and 2,107 controls. The samples used from the three European GWAS were independent: the main GWAS publication used Identity by descent (IBD) analysis in PLINK 1.9b (www.cog-genomics.org/plink/1.9/) (27) to remove individuals from Genentech with IBD > 0.125, we used these data and applied the same analysis to the SLEGEN data.

Chinese samples were from previously published GWAS from Anhui (1,047 cases and 1,205 controls) (28) and Hong Kong (612 cases and 2,193 controls) (29, 30).

Clinical sub-phenotypes were available for the SLE main cohort and SLEGEN cohort, which were documented according to the standard American College of Rheumatology (ACR) classification criteria. Subgroups of patients with renal disease or without renal disease were identified according to the sub-phenotype data using ACR classification. Following quality control, the sample size of patients with renal disease, lupus nephritis (LN+) were 1,152 and 146; while patients without renal disease (LN-) were 1,949 and 378 in the SLE main cohort and SLEGEN cohort, respectively. More details are presented in **Table S1**.

## Genome-wide association study (GWAS)

### SLE GWAS

SLE GWASs were performed in genotyped SNPs including principal components consistent with the original publications in all three independent cohorts.

### SLE Renal GWAS within SLE cases

The SLE Renal GWASs were performed within SLE cases, i.e., genome-wide associations of patients with renal disease (SLE Renal+, cases) and patients without renal disease (SLE Renal-, controls) in two independent cohorts, i.e., the SLE main cohort and the SLEGEN cohort. For Renal GWASs, we pre-phased the genotyped data using the SHAPEIT algorithm (31) and then used IMPUTE2 (32) to impute to the density of the 1000 Genome reference data (phase 3 integrated set, release 20130502) (33) (data unpublished). All case-control analysis was carried out using the SNPTEST algorithm (34). SNPs with imputation INFO scores of < 0.7 and MAF (minor allele frequency) < 0.001 were removed. After quality control (QC), there were 21,431,070 SNPs left for further analysis. Moreover, a genome-wide association meta-analysis of the SLE main cohort and SLEGEN cohort was performed using the summary statistics derived from the two Renal GWASs. A standard threshold of *P* ≤ 5e-08 was used to report genome-wide significance and a *P* ≤ 1e-05 was used to report suggestive associated signals.

### Polygenic analysis

We tested for non-zero standardized effect sizes (Z scores) for SLE association in the Genentech data for groups of SNPs stratified by their *P* values in the SLE main cohort. The Z scores in the Genentech data were polarized with respect to the SLE main cohort in that the effect allele was set to be the risk allele in the SLE main cohort. Under the null hypothesis the Z scores will have zero mean, while under the alternative the mean will be positive. SNPs were stratified by *P* value intervals of 1-0.9, 0.9-0.8, 0.8-0.7, 0.7-0.6, 0.6-0.5, 0.5-0.4, 0.4-0.3, 0.3-0.2, 0.2-0.1, 0.1-0.00. We would expect a positive mean for SNPs with very small *P* values in the main SLE cohort as these will be enriched for true positives, while the same is not necessarily true over other *P* values ranges unless there are more widespread true associations with very weak effects. We also ran this analysis on renal association standardized effect sizes (Z scores) again polarized with respect to SLE association and stratified by SLE *P* values. In all analyses, we used an LD clumped set of SNPs with an R^2^ threshold of 0.1. When comparing the SLE main cohort to the Genentech cohort or the SLEGEN cohort, we limited the clumping to SNPs that overlap the GWASs.

### Genetic risk score derivation

A Genetic risk score (GRS) is a quantitative trait of an individual’s inherited risk based on the cumulative impact of many genetic variants, which is calculated according to the method described by Hughes et al (35), taking the number of risk alleles (i.e., 0, 1 or 2) for a given SNP and multiplying this by its corresponding estimated effect - β coefficient, i.e. the natural log of its odds ratio (OR). The cumulative risk score in each subject was calculated by summing the risk scores from the target risk loci:

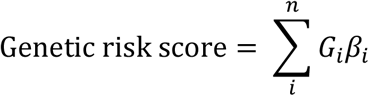

where *n* represents the number of SLE risk loci, *G_i_* is the number of risk alleles at a given SNP, and *β_i_* is the effect size of the risk SNP *i*.

We used two approaches to select SNPs for GRS calculation. The first approach – a weighted GRS was derived from all published independent SLE risk SNPs (**Table S2**) – including 78 SLE susceptibility loci (without the X chromosome), consisting of 93 SNPs outside of the MHC region and 2 independent tag SNPs in the MHC region for two SLE associated HLA haplotypes. The risk allele for each SNP is derived from its original publication, which is summarized in a recent review (11) and the effect size used in the GRS was generated from each GWAS used as a training set. Each GRS for four SLE cohorts (24, 25, 28–30) was generated using R version 3.4.3.

The second approach – LD clumping and thresholding – was used to build 32 GRSs. Clumping and thresholding scores were built using a *P* value and linkage disequilibrium (LD)-driven clumping threshold in PLINK version 1.90b (www.cog-genomics.org/plink/1.9/) (27). In brief, the algorithm forms clumps around SNPs with association *P* values less than a provided threshold (Index SNPs). Each clump contains all SNPs within a specified window of the index SNP that are also in LD with the index SNP as determined by a provided pairwise correlation threshold (r^2^) in the training data. The algorithm loops through all index SNPs, beginning with the smallest *P* value and only allowing each SNP to appear in one clump. The final output should contain the most significant disease-associated SNP for each LD-based clump across the genome. Note that when performing LD clumping, we firstly removed the X-chromosome and the MHC extended region (24-36MB) and kept all other autosomal SNPs. Then we included the MHC region by using two tag SNPs for two well-known HLA haplotypes in SLE, i.e. rs2187668 for HLA-DRB1*03:01 and rs9267992 for HLA-DRB1*15:01 for the European cohort and rs9271366 for HLA-DRB1/HLA-DQA1 and rs9275328 for HLA-DQB1/HLA-DQA2 for the Chinese cohort (**Table S2**). A GRS was built using the genotypes for the index SNPs weighted by the estimated effect sizes (β). Specifically, when training the GRS in the SLE main cohort and testing in the SLEGEN cohort, we performed a GWAS on the genotyped SNPs in the SLE main cohort and generated 32 lists of clumped SNPs over a set of P values (*--clump-p1:* 0.1, 0.01, 1e-03, 1e-04, 1e-05, 1e-06, 1e-07,and 5e-08), r^2^ *(--clump-r2:* 0.2 and 0,5) and clumping radius *(--clump-kb:* 250 and 1000). The 32 lists of SNPs were then used to generate 32 GRSs by summing across all variants weighted by their respective effect size for samples in the SLEGEN cohort. We performed this analysis using all three cohorts in European population with one dataset as training and the other as a test set, generating six training-and-testing pairs. We also performed a cross population analysis between European and Chinese populations.

### Receiver Operating Characteristic (ROC) curves for model evaluation

The GRS with the best discriminative capacity was determined based on the maximal Area under the ROC curve (AUC) with SLE or RENAL as the outcome and the candidate GRS as the predictor. AUC confidence intervals were calculated using the ‘*pROC* package within R and the difference between the ROC curves was determined with the ‘*roc.test*’ function, which used a non-parametric approach, as described by De Long et al (36). To assess the degree to which the age of SLE onset contributes to the prediction of renal involvement within SLE cases, we generated ROCs as above with the GRS and compared to ROC curves with SLE age onset as a single predictor and the ROC with both GRS and age onset as predictor(s).

### Partitioning the genetic risk of renal disease

Since a continuous score is difficult to interpret on an individual level when a physician needs to explain the results of the GRS to a patient, we partitioned SLE patients into quintile according to genetic dosage (SLE GRS). We used a chi-square test to study the association of the partitioned GRS and renal risk. The odds ratios of renal risk were then calculated compared to the reference group - the first quintile GRS group.

To test whether the GRS correlated with renal disease independently of age-of-onset, we partitioned SLE patients into two groups according to their age of onset, with a cutoff at age of 30 - patients with age above 30 were defined as ‘Late age onset’ and others as ‘Early age onset’. A two-way ANOVA test was then performed with the function *‘aoV* in R, with *aov*(*GRS ~ age group * renal group*). All statistical analyses were conducted using R version 3.4.3 software (https://www.r-project.org/).

## Results

### The best GRS in SLE prediction

Among the GRSs generated from LD clumping and thresholding, the predictor with the best discriminative capacity was the one derived from SNPs clumping at *P* threshold (*P_th_*) of 1e-05 with R^2^ < 0.2 in the SLE main cohort and tested in both the SLEGEN (AUC = 0.72; 95% C.I. = 0.69-0.74) and Genentech (AUC = 0.67; 95% C.I. = 0.66-0.69) cohorts (**Figure 1** & **Table S3**), suggesting there may be more true positive signals than the genome-wide significant ones involved in the risk of SLE. This performance was not due to population structure as the GRS added significantly more (*P* = 2.2e-16 and *P* = 7.78e-14) to the AUC than principal components in both Genentech and SLEGEN respectively. In fact, the predictive performance of the GRS using all pairs of training and test data was maximised using SNPs below the standard genome-wide threshold (**Table S3**). This evidence for polygenicity was also seen in an analysis of the association statistics (*Z* scores) in the Genentech GWAS polarised to the risk allele in the main GWAS, partitioned by their association *P* value in the main GWAS (see **Methods**). Here, we found evidence (**Figure 2** & **Table S4**) against a zero mean (*P* = 3.91e-04) for the Z scores in Genentech data for SNPs with *P* values between 0.3 and 0.2 in the main GWAS.

**Figure 1.**
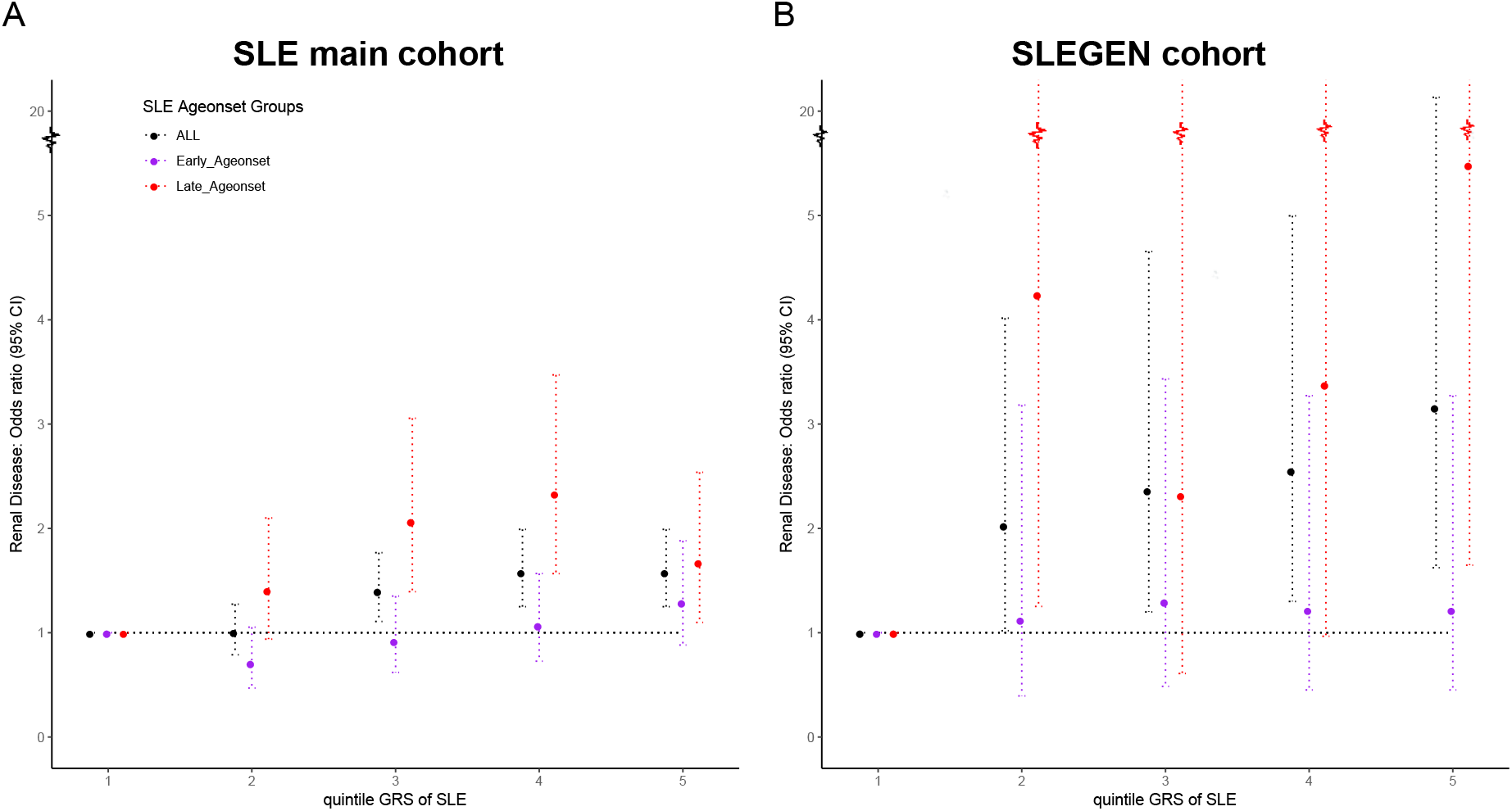
ROCs and AUCs of models in SLE prediction in European cohorts and between ancestries. GRSs for the prediction of SLE in the SLEGEN cohort (**A**) and Genentech cohort (**B**) were generated from SNPs of LD clumping and threshold derived from the SLE main cohort. All GRSs for the training-and-validation in European cohorts were generated with two MHC tag SNPs derived from the European GWAS (See **Methods**). GRSs for the prediction of SLE across populations (**C**) and (**D**) were generated from SNPs of LD clumping and threshold without MHC tag SNPs. The ‘GRS at *P_th_* represented the GRS in the SLE prediction model, which was derived from the LD clumping at the according GWAS P value threshold.

We found that the genetic risk score trained in our European (EUR) data predicted SLE in the Chinese (CHN) data well (**Figure 1C & 1D**) with an AUC (0.64) when using the best approach for GRS in the Europeans (R^2^ < 0.2 for all SNP pairs and using SNPs that passed the *P* value threshold of 1e-05). The range of AUC values over all *P* value thresholds for SNP inclusion was [0.60 – 0.64]. The results when training in the CHN and predicting in EUR were similar: AUC = 0.64 when using the best approach for GRS in the Europeans (R^2^ < 0.2 for all SNP pairs and using SNPs that passed the *P* value threshold of 1e-05) and range of AUC values over all *P* value thresholds for SNP inclusion was [0.55 – 0.64].

### Lupus Nephritis GWAS within SLE cases

Lupus Nephritis (LN) occurs in approximately half of all SLE patients, and its frequency ranges from 25% to 75% depending on the population studied (37). About one third of European SLE patients experience renal disease (38). Until recently, one of the most common causes of death in SLE patients was kidney failure. According to the lupus severity index (LSI) using the ACR criteria developed by Bello et al (39), renal involvement has the highest impact and particular strongly associated with disease severity, hence we chose LN as a proxy of SLE severity in this study.

The within case LN GWAS in the SLE main cohort, which comprised 1152 SLE patients with renal disease (LN+) and 1949 patients without renal disease (LN-), did not identify any genome-wide significant associated loci (*P* ≤ 5e-08) (**Figure S1A**). Consistently, no inflation (genomic inflation factor: λ = 1.014) was observed in the QQ plot (**Figure S1D**). Similarly, none of the SNPs reached genome-wide significance in the SLEGEN cohort (25) (λ = 1.023) (**Figure S1B & 1E**). In addition, no variant passed genome-wide significance in the meta-analysis of the SLE main cohort and SLEGEN cohort for Renal GWAS (λ = 0.9565) (**Figure S1C & S1F**). Summary association statistics for SNPs with *P* ≤ 1e-05 are provided in **Table S5 and S6**.

We did, however, see evidence that SNPs with very strong evidence for association with SLE (*P* ≤ 1e-05) were associated with LN. This was evident from an analysis of the renal association statistics (Z scores) polarised to the risk allele for SLE. There was strong evidence (**Figure 2 & Table S4**, *P* = 8.72e-08) against a zero mean for the Renal Z scores for SNPs with *P* ≤ 1e-05 for SLE in the main cohort. This result was replicated in the SLEGEN study with *P* = 2.42e-03 (**Figure 2 & Table S4**). The finding of renal association with SNPs showing very strong evidence for association with SLE could be exploited for prediction of disease progression and we explore this below.

**Figure 2.**
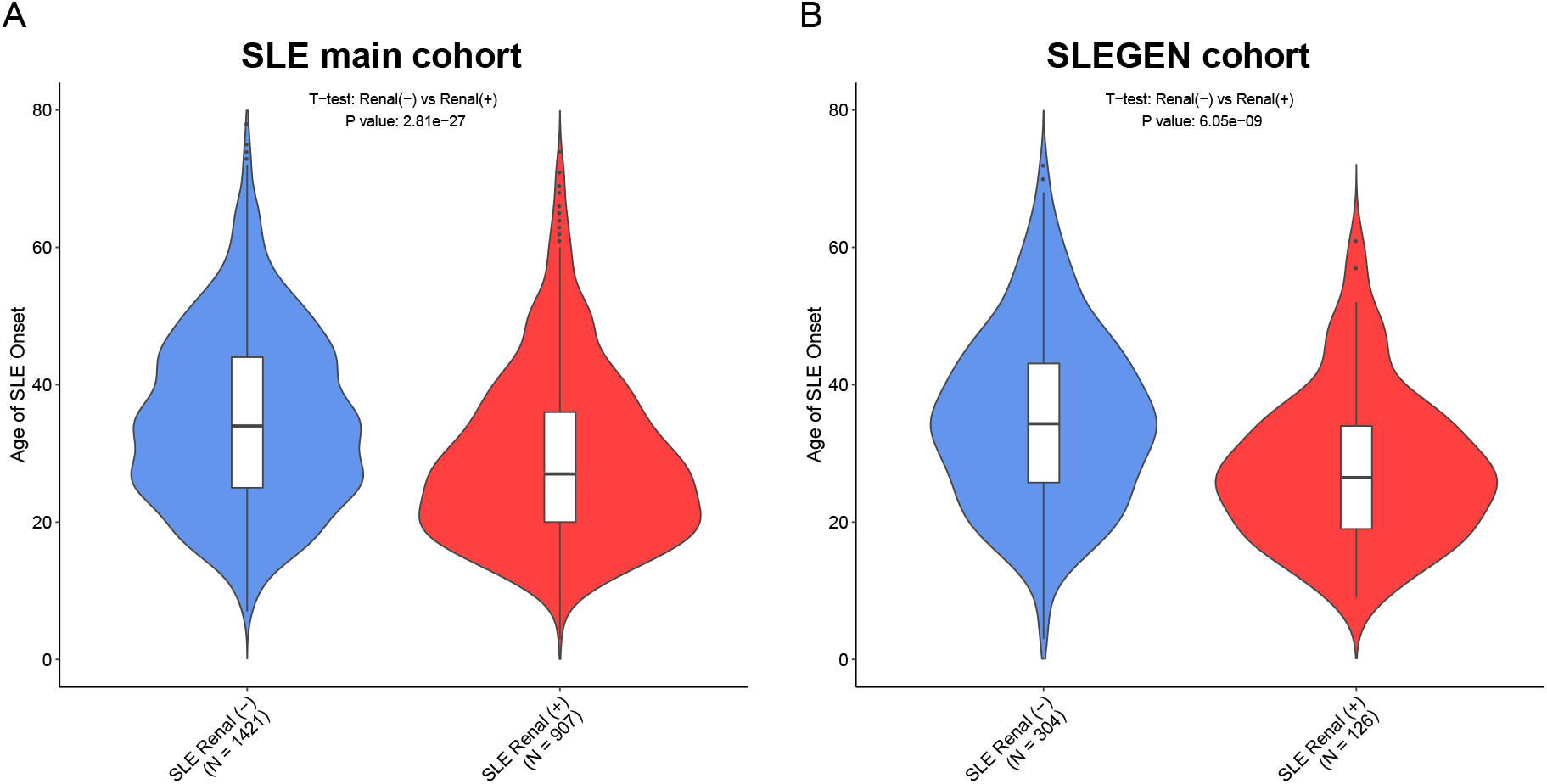
Polygenic test of SLE and Renal disease. Polygenic test of SLE in Genentech cohort (**A & B**) and polygenic test of Renal disease in the SLE main cohort (**C & D**) and SLEGEN cohort (**E & F**). The SLE main cohort was used to generate a *P* value for each SNP, to stratify the SNPs into groups for the Z score calculation of SLE association or Renal association.

### Genetic risk loading of SLE is significantly higher in LN+ patients

While we observed that no individual SNPs were significantly associated with renal involvement in the SLE cases, we did show that there was a deviation from zero mean for renal Z scores taken from SNPs with very strong evidence for association with SLE. In view of this finding, we investigated the correlation between the SLE GRS and renal disease in all SLE cases. To accomplish this, we used the GRS derived from a list of published SLE associated SNPs (11) for the comparison of the SLE genetic risk burden in patients with and without renal disease. As expected, the GRS was higher in the SLE patients compared to healthy controls in both independent cohorts (**Figure 3**).

**Figure 3.**
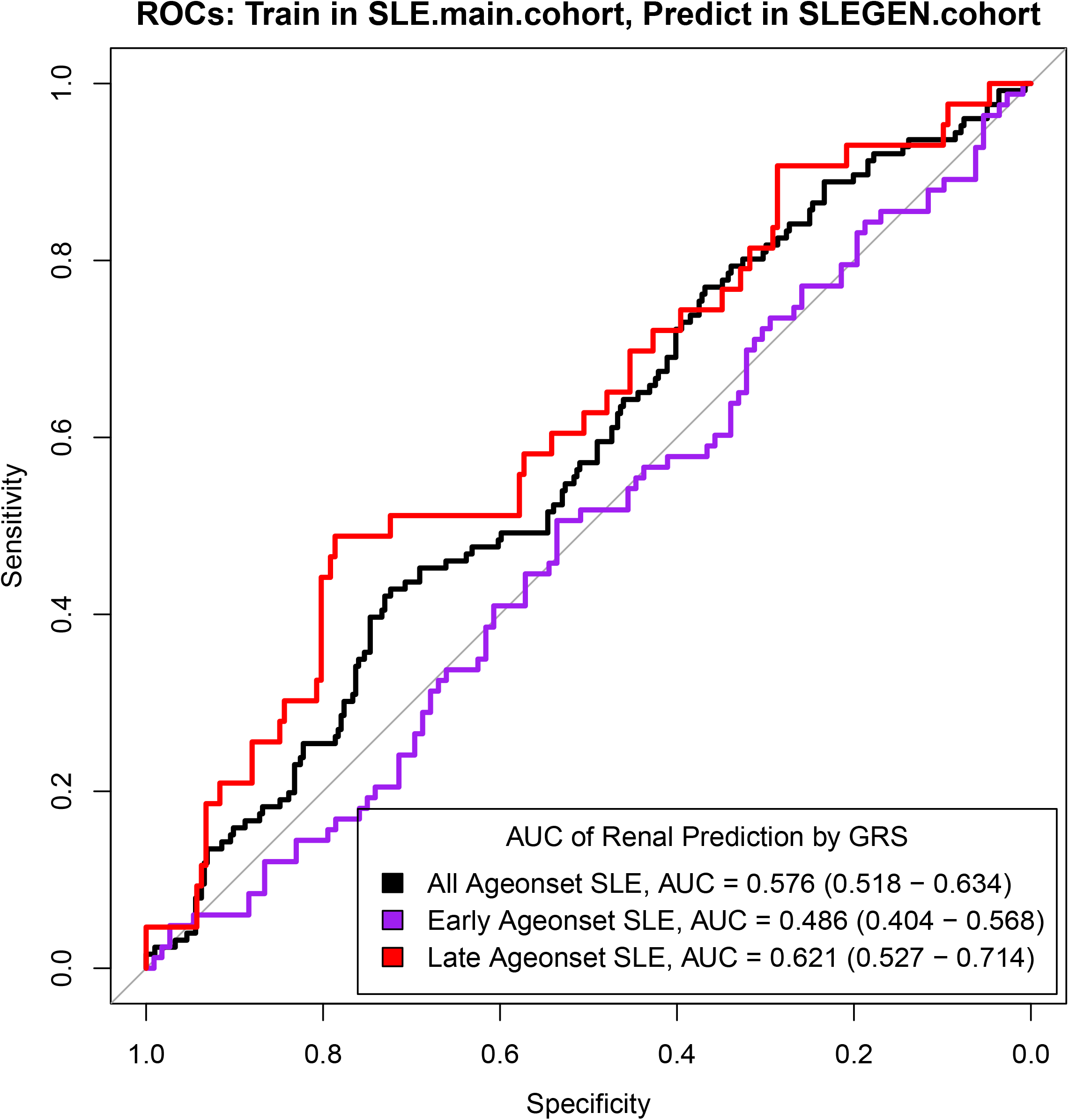
GRS over levels of disease: Controls / SLE Renal (-) / SLE Renal (+). The violin-and-box plots show the summary GRS for each level of the disease in the SLE main cohort (**A**) and the SLEGEN cohort (**B**). The violins show the distribution of the GRS across each group. The bottom line of the box inside the violin is the 1st quantile, the top line is the 3rd quantile, and the box is divided at the median. Sample size (N) of each group is showed within brackets below the group name. Note that GRS for SLE main cohort and SLEGEN cohort are generated by 93 non-MHC SNPs and 2 MHC tag SNPs - a total of 95 SNPs (**Table S2**).

A significantly higher GRS was observed in the group of patients with renal disease (LN+) compared to patients without renal disease (LN-) (**Figure 3**). In the SLE main cohort, the mean (SD) of the GRS was 18.1 (1.64) for LN+ patients and 17.8 (1.65) for LN-patients (*P* = 1.60e-07); the mean (SD) for the SLEGEN cohort was 18.2 (1.66) for LN+ patients and 17.6 (1.69) for LN-patients (*P =* 0.0010). Moreover, we saw a significant increasing trend of GRS over levels of diseases: Healthy control, LN-patients, and LN+ patients, in the SLE main cohort and the SLEGEN cohort (**Figure 3**).

### Genetic risk of nephritis and age of onset in SLE

We partitioned the SLE cases into five groups according to quintiles for GRS to show the risk of renal involvement. We observed over 1.5 folds higher risk of renal disease (OR = 1.58; 95% C.I. = 1.25-1.99; *P* = 0.00015) between the top and bottom quintiles of GRS in the SLE main cohort (**Figure 4A**). This is replicated in the SLEGEN cohort (**Figure 4B**), with odds ratios of 3.16 (95% C.I. = 1.62-6.13; *P* = 0.00091). A significantly earlier age of SLE onset was observed in those with renal disease compared to those without renal disease. In the main cohort (**Figure 5A**), the mean (SD) for age of disease onset was 29yrs (12) for LN+ patients and 35yrs (13) for LN-patients (*P* = 2.8e-27); the means for the SLEGEN cohort (**Figure 5B**) were 28yrs (11) and 35yrs (13) for LN+ and LN-, respectively (*P* = 6.05e-09). When testing the association of GRS with age of onset in the SLE main cohort, a significant correlation was present – the higher the GRS, the earlier age of SLE onset (*P* = 4.59e-12). This correlation was also detected in the SLEGEN cohort (*P* = 0.021) and the combined Chinese cohort (*P* = 1.57e-06).

**Figure 4.**
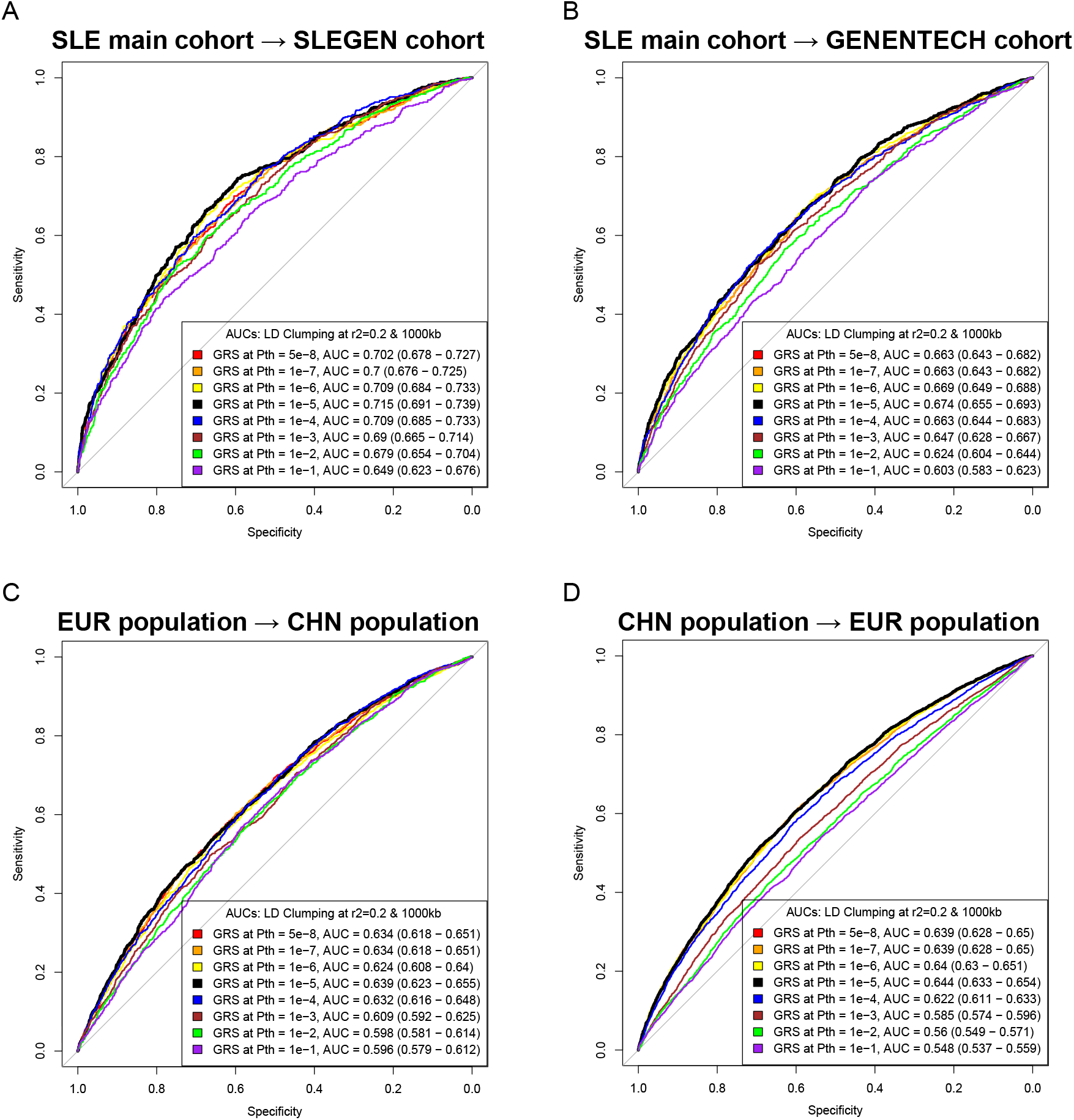
Relationship of quintiles of the GRS and risk of renal disease within SLE patients. Plots show the odds ratios of Renal disease for the SLE main cohort (**A**) and the SLEGEN cohort (**B**), comparing each of the upper four GRS quintiles with the lowest quintile; dotted lines represent the 95% confidence intervals (C.I.); horizontal black dotted lines represent OR = 1.

**Figure 5.**
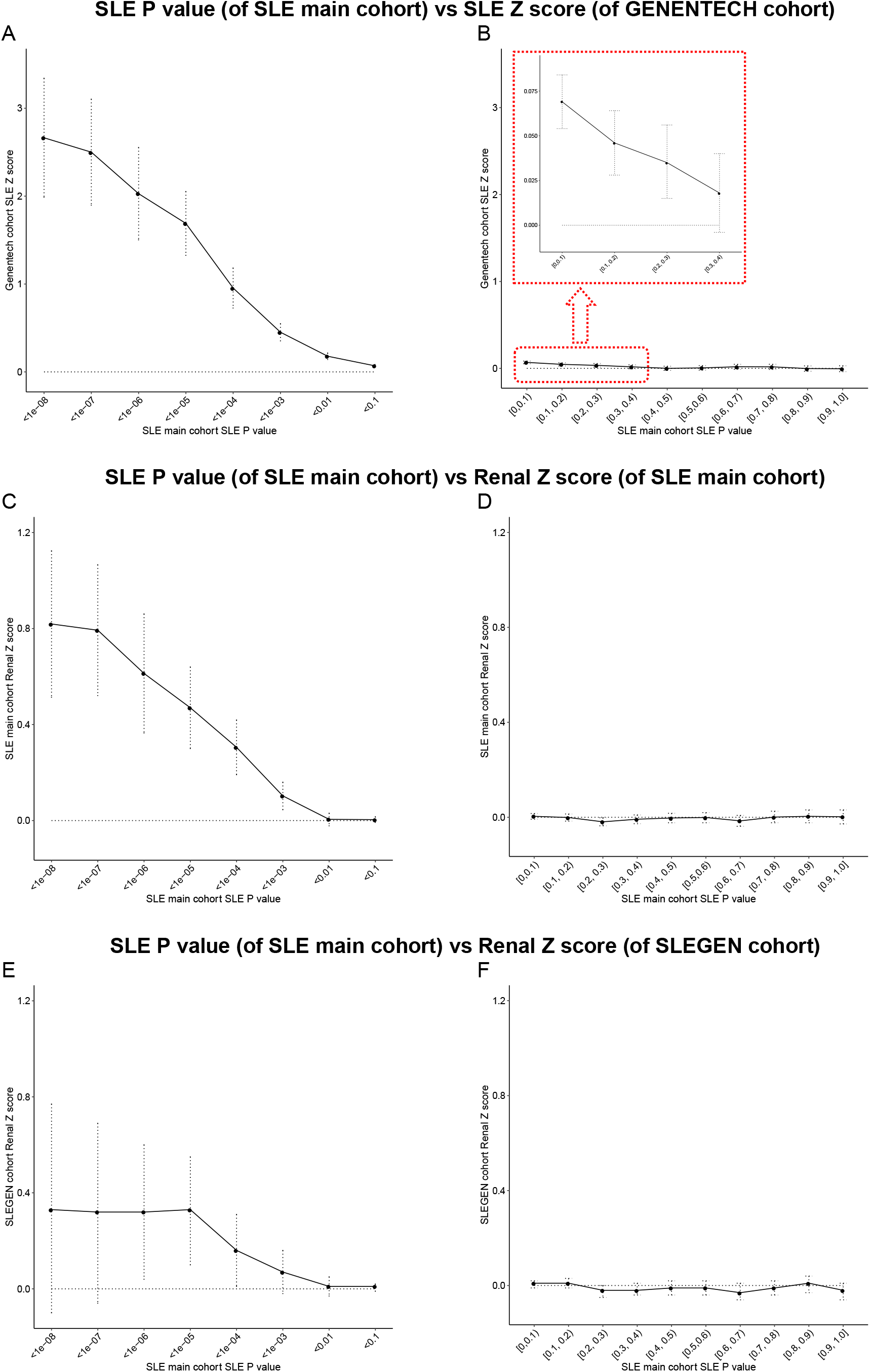
Age of SLE onset in patients of Renal(-) / Renal(+). The violin-and-box plots show the age of SLE onset for each level of the disease in the SLE main cohort (**A**) and the SLEGEN cohort (**B**). The violins show the distribution of the Age of SLE onset across each group. The bottom line of the box inside the violin is the 1st quantile, the top line is the 3rd quantile, and the box is divided at the median. Sample size (N) of each group is showed within brackets below the group name.

To test whether the GRS correlated with renal disease independently of age-of-onset, we partitioned SLE patients into two groups according to their age of onset, i.e. ‘Late age onset’ and ‘Early age onset’ and performed a two-way ANOVA test (See Methods). The GRS was shown to positively correlate with both renal disease and early age-of-onset (*P*_Renal_ = 7.64e-05 and *P*_age-of-onset_ = 1.06e-09) in the SLE main cohort, with significant association with renal disease in the SLEGEN cohort but marginal evidence for age-of-onset (*P*_Renal_ = 0.0288 and *P*_age-of-onset_ = 0.0513), while we found that there was no statistically significant interaction between renal and early age-of-onset in the SLE main cohort (*P*_Interaction_ = 0.795) and marginal evidence in the SLEGEN cohort (*P*_Interaction_ = 0.0511) (**Figure S2**). Notably, we found that GRS was a better predictor of renal disease in the ‘Late age onset’ group (AUC = 0.62) compared with the ‘Early age onset’ group (**Figure 6**).

**Figure 6.**
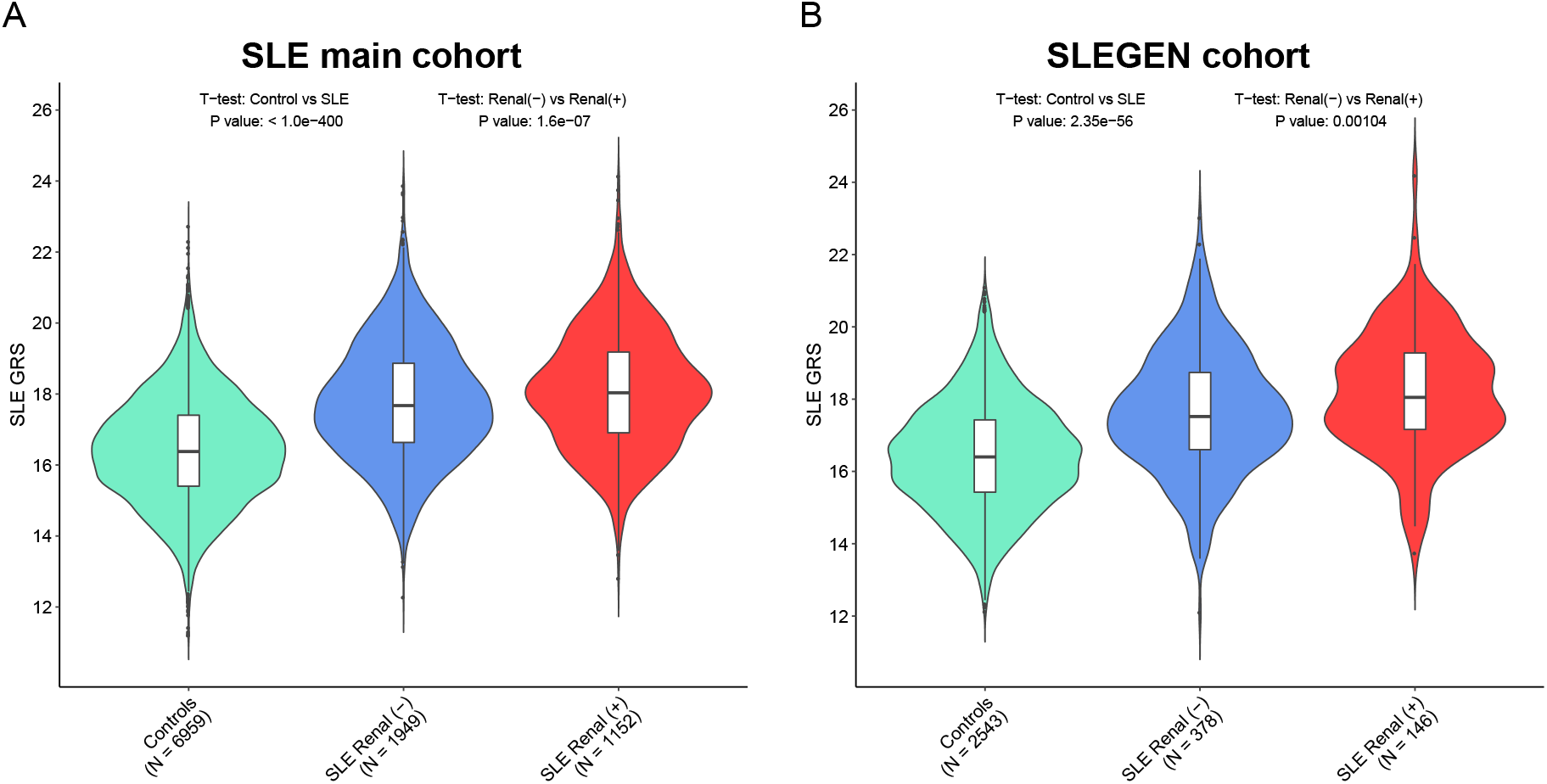
ROC Curves for models predicting a diagnosis of Renal disease in SLE patients using GRS, split by age-of-onset. The models were trained in the SLE main cohort and tested in the SLEGEN cohort. The plots showed the ROC curves in the prediction of renal disease in SLE patients with GRS as a predictor, The ROC curve in black was trained and tested with all SLE samples, the purple curve was trained and tested in the ‘Early age onset’ patients (≤ 30yrs), and the red curve was trained and tested in the ‘Late age onset’ group. AUC, area under the ROC curve is showed with 95% C.I. in brackets.

Finally, we assessed the predictive ability of the partitioned SLE GRS (quintile GRS, see methods) over the two age-of-onset groups. In the main SLE cohort there is a clear and significant risk effect for renal involvement with increasing GRS in the ‘Late age of onset’ group, but no significant effect in the early onset group. We observed over two folds higher risk of renal disease (OR = 2.33; 95% C.I. = 1.57-3.47; *P* = 3.76-05) between the upper fourth quintile and the bottom quintile in the ‘Late age onset’ group in the SLE main cohort (**Figure 4A**). The results were similar in the SLEGEN cohort, with the risk of renal disease between the top and bottom quintile of GRS being over five times (OR = 5.48; 95% C.I. = 1.65-18.3; *P* = 0.00664) (**Figure 4B & Table S7**) in patients of ‘Late age onset’ but no significant differences in those with ‘Early age onset’. These results are robust to the chosen threshold in the definition of ‘Late age onset’ and ‘Early age onset’ (**Table S7**)

## Discussion

GRS has been showed to be predictive for several diseases including cardiovascular disease (AUC = 0.81,95% C.I. = 0.81-0.81) (12), inflammatory bowel disease (AUC = 0.63, 95% C.I. = 0.62–0.64) (12) and breast cancer (AUC = 0.63, 95% C.I. = 0.63-0.65) (40). However, in many of these applications the AUC values are dependent on inclusion of age and sex for prediction and so the AUC due to genetics alone would have been substantially lower (41). We have shown that a SLE GRS using only SNPs has good predictive power with AUC approaching 0.7 over a range of settings when trained and tested between three European GWAS. We also used two combined Chinese studies’ data as both independent validation and a test of cross populations prediction performance. In both populations we show that, when using GWAS data as a training set, a GRS using SNPs with association *P* values well below genome-wide levels of significance has the best predictive performance. This, along with other studies that have reinvestigated SLE GWAS data (42), is further evidence that SLE is a polygenic disease with many risk variants as yet undiscovered, and that more powerful studies could lead to useful predictive models. Genetic risk scores may also have utility in prediction of disease severity and we find evidence for this to be so for SLE. Our data show that renal involvement is not related to specific genetic factors or particular genes but simply to genetic load of risk alleles.

Until recently, the most common cause of death in SLE patients was kidney failure. Though the frequency of death from kidney disease has decreased sharply due to better therapies *(e.g.* dialysis and kidney transplantation), kidney failure is still potentially fatal in some people with SLE and causes significant morbidity. According to the lupus severity index (LSI) using the ACR criteria developed by Bello et al (39), renal involvement had the highest impact and particularly more strongly associated with disease severity, hence we used renal involvement as a proxy of SLE severity in this study. In the SLE within-case renal GWASs, we observed no genome-wide significant signals in either the SLE main cohort or the SLEGEN cohort, or metaanalysis of these two. Both datasets had genetic variants with less stringent *P* values (*P* ≤ 1e-05) for renal association, but none of them were replicated in the other cohort. Considering the sample size of both cohorts are relatively small, we applied an online genetic power calculator (http://zzz.bwh.harvard.edu/gpc/) to calculate the power of our current sample size for the GWAS study (**Table S8**). We assumed the effect sizes of SLE renal risk alleles is similar to that seen in SLE GWAS, so the odds ratio (OR) of the risk allele would be between 1.0 and 2.0. Therefore, we calculated power under a variety of parameters, including OR, risk allele frequency (RAF) and alpha. As showed in **Table S8**, we have a power of ≥ 0.8 to detect a genetic risk variant with an OR = 1.4 and RAF = 0.3 or an OR = 1.5 and RAF = 0.2 when alpha = 5e-08. However, if we assume the renal associated variants are as weak as most of the SLE associated variants (OR < 1.2), then we are under powered (< 0.8) to detect the true renal associations at the GWAS significant threshold of *P* = 5e-08 in the current study.

We did however find evidence that SNPs most associated with SLE (*P* < 1e-05) were enriched for associations with SLE renal involvement. Specifically, the renal association *P* values of the 95 SNPs (of 77 published SLE risk loci) in the SLE main cohort and the SLEGEN cohort are strongly inflated as shown in the QQ plots (**Figure S3**), suggesting the cumulative genetic burden from multiple SLE risk genes with modest effect. So we then tested the hypothesis that the genetic risk loading of SLE may correlate with kidney involvement. Therefore, a genetic risk score (GRS) using published SNPs with robust evidence for association with SLE was derived for the prediction of SLE renal disease. In both European cohorts, the SLE main cohort and the SLEGEN cohort, the GRS was significantly higher in patients with renal disease than patients without. In addition, patients with a higher GRS were more likely to have renal involvement at a younger age, indicating the strong genetic background of SLE development. These findings provide more evidence to support the opinion that younger-age onset lupus is generally more severe than older-onset lupus as reported previously (43–45).

Our analysis of Renal disease in SLE patients has shown that, while we find no SNPs significantly associated with renal disease, the fact that SLE associated variants correlated with renal using a GRS suggests that many SLE associated variants are also risk for renal involvement albeit with likely weaker effects (Odds ratios). We find that the GRS and age-of-onset are correlated but the GRS is associated with renal involvement independently of age-of-onset with no interaction observed. The GRS performs better for predicting renal disease in patients with late age-of-onset. We also find that a stratified GRS may be a more viable option for predicting renal disease, where we estimate significantly high relative risks for those in the tails of the GRS distribution in both of our European studies that had renal data.

A limitation of this study is that we were not able to replicate our renal results in the Chinese as renal data were not available. Renal involvement in Chinese is more common than in Europeans; the Chinese SLE patients are more heterogeneous, suffer from more severe clinical manifestations and earlier age of onset. The use of GRS for predicting SLE severity in Chinese may not have the same utility as in Europeans where we find the stronger association in the late onset patients. Nevertheless, our results in Chinese showing a correlation between age of onset and SLE GRS suggest that in this population disease severity is also driven by load of disease associated variants.

This is the first study to investigate accumulated genetic risk and its relationship with the susceptibility and severity of SLE with data in Chinese and European populations. We found that the higher the GRS, the younger onset of SLE in both populations. Within the European population and across the Chinese and European populations we find that a genetic risk score incorporating LD pruned SNPs (at R^2^ = 0.2) with modest (*P* < 1e-05) evidence for association with disease predicts SLE with AUC of 0.64 and above. In the European data we see that in patients of late onset, a higher GRS means patients are more likely to suffer from more severe disease. In brief, age of onset incorporating a GRS may assist early prediction of lupus nephritis in a clinical setting. Nevertheless, more clinical studies and multi population data are needed to validate the usefulness of this application.

## Acknowledgements

We thank Dr. Amy W.L Bulter and Mr. Akmal Droubi for excellent clinical data administration on both SLE cohorts. We thank Professor Cathryn Lewis for critical review of this article.

