## Supplementary material Figures for "Genome-wide assessment of genetic risk for systemic lupus erythematosus and disease severity"

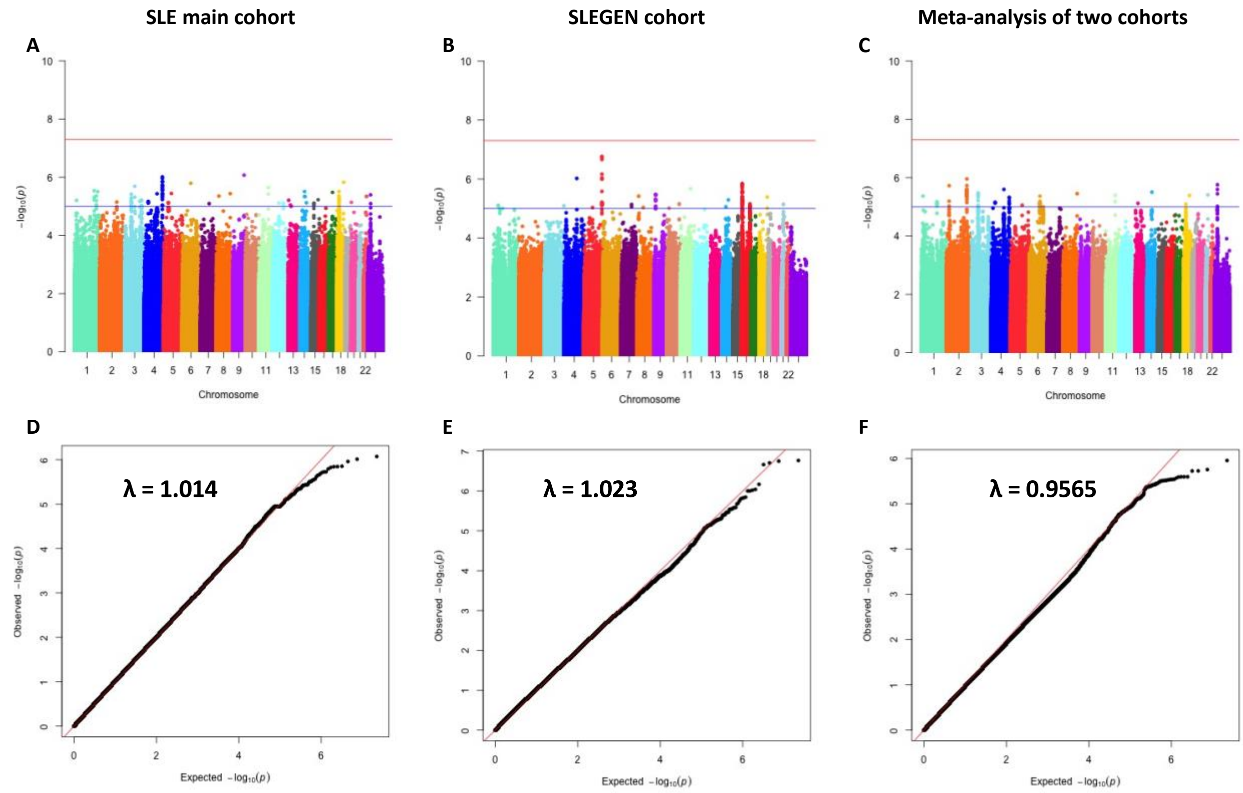


**Figure S1. Genome-wide scans of LN associated variants.**

(Upper) Manhattan plots showing the -log10-transformed p values (y axis) against physical genomic position (x axis) for each SNP in the SLE main cohort (**A**), the SLEGEN cohort (**B**), and the meta-analysis of these two cohorts (**C**). The red horizontal line represents the threshold for genome-wide significance (P ≤ 5e-08) and the blue horizontal line represents the threshold for suggestive significance (P ≤ 1e-05).

(Lower) Quantile-quantile plots showing the observed distribution of -log10-transformed p values (y axis) by the expected distribution (x axis) under the null hypothesis of no association (diagonal line) for the SLE main cohort (genomic inflation factor, λ = 1.014) (**D**), the SLEGEN cohort (λ = 1.023) (**E**), and the meta-analysis of these two cohorts (λ = 0.9565) (**F**).

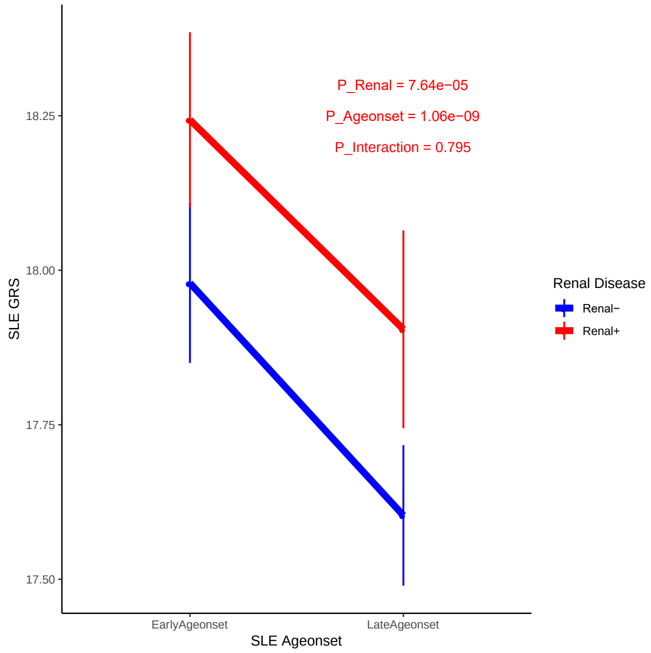


**Figure S2. Relationship of GRS and age onset in Renal disease.**

The age of SLE onset ≤ 30 years was defined as “Early onset” and > 30 years was defined as “Late onset”. For each age onset and renal group, the GRS was plotted with mean and 95% CI for the SLE main cohort.

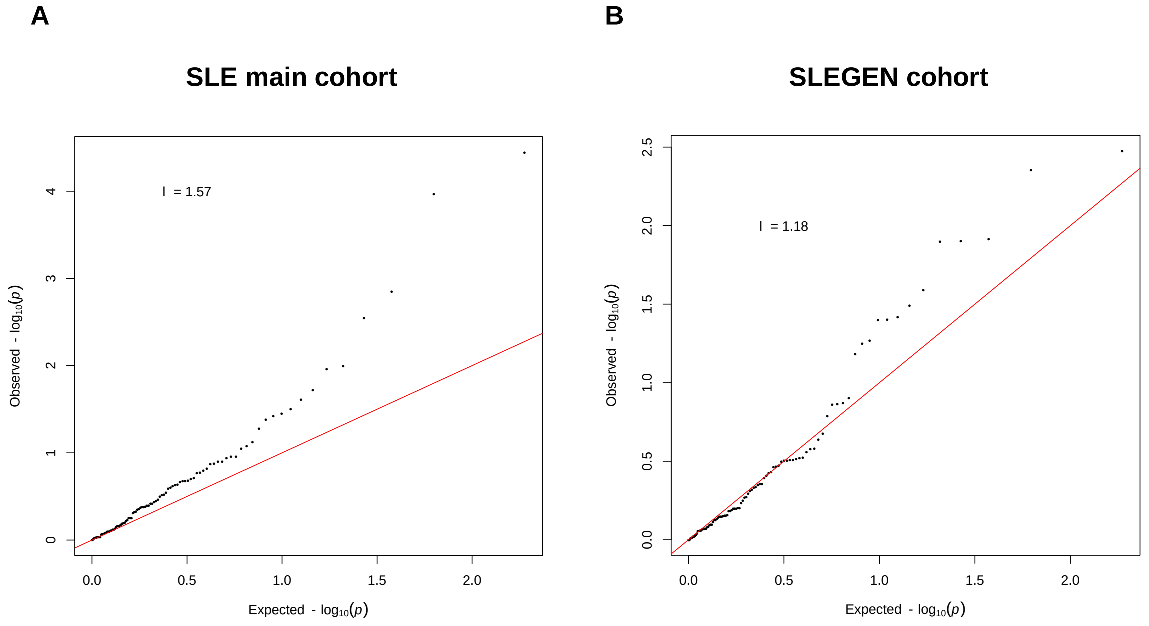


**Figure S3. Quantile-quantile plots of Renal association results**

QQ plots showed the observed distribution of -log10-transformed p values (y axis) by the expected distribution (x axis) under the null hypothesis of no association (diagonal line) for the SLE main cohort (**A**) and the SLEGEN cohort (**B**). The *P* values for the QQ plots were derived from Renal association test of the 95 SNPs (**Table S3**) which used for the GRS calculation.
